# The role of falx cerebri in the selective vulnerability of the splenium in the corpus callosum

**DOI:** 10.64898/2026.05.31.729036

**Authors:** Zhou Zhou, Svein Kleiven

## Abstract

The corpus callosum, the largest white matter tract connecting the cerebral hemispheres, is anatomically divided into the genu, midbody, and splenium. Among these subregions, the splenium is particularly vulnerable to traumatic injury, with the underlying biomechanics largely unknown. We hypothesized that the falx cerebri contributes to splenial vulnerability because of its anatomical proximity to the posterior corpus callosum and its substantially greater stiffness relative to brain tissue. To test this hypothesis, a high-resolution finite element head model with explicit representations of the corpus callosum subregions was used to simulate ten head impacts with and without an anatomically and mechanically detailed falx. Peak strain, strain rate, and shear stress were quantified and compared using linear mixed-effects models. The results showed the inclusion of the falx consistently increased strain, strain rate, and shear stress in the splenium relative to the genu and midbody, whereas this preferential splenial response was not consistently observed without the falx. Statistical analyses confirmed significant region-dependent effects, with falx-induced increases significantly greater in the splenium than in the other subregions (*p* < 0.05). These findings verified that the falx selectively amplifies mechanical loading within the splenium, providing a biomechanical explanation for its frequent involvement in neurotrauma patients.

## Introduction

Traumatic brain injury (TBI) is a significant public health problem. In Europe, the annual number of TBI-related victims is estimated to be 2.5 million, of which 1 million require hospital admission and 75,000 die [1]. In Sweden, falls are the leading cause of TBI, with incidence rates showing only a minor decrease in men (6%) and no significant reduction in women from 2008 to 2022 [2]. In France, approximately 25% of traffic-related injuries involve the head region, among which mild TBI without loss of consciousness is the most common (59.7%) [3]. TBI is the leading cause of death in young adults, and its incidence rate in the elderly is alarmingly increasing [4]. Uncovering the brain injury mechanisms and developing advanced head protective gear are central to mitigating this TBI-related epidemic.

As the largest white matter structure connecting the two cerebral hemispheres, the corpus callosum attracts significant interest in neurotrauma research. Early autopsy examinations regarded corpus callosum injury as a hallmark pathology for diagnosing and grading axonal injury [5-7]. At the mild injury level, corpus callosum damage has been associated with impairments in perception, speech, orientation, and reaction performance [8, 9]. Advanced imaging techniques, such as diffusion tensor imaging (DTI), are highly sensitive to white matter compromise within the central tracts of the corpus callosum [10-12]. DTI-based parameters, such as fractional anisotropy and mean diffusivity, in the corpus callosum have been proposed as non-invasive biomarkers for underlying structural integrity and pathological alteration [13, 14]. Of particular relevance to the current study, comparable interests are noted from the biomechanical community to study the mechanical responses of the corpus callosum during head impacts. For example, Kleiven [15] developed a three-dimensional (3D) finite element (FE) head model with detailed descriptions of brain anatomy, material, and interface to calculate the maximum principal strain in the corpus callosum, a metric which was later identified by Hernandez *et al*. [16] as the strongest predictor of concussion out of eighteen candidates. Similarly, McAllister *et al*. [17] reported that the maximum principal strain and strain rate significantly correlated with changes in fractional anisotropy and mean diffusivity within the corpus callosum of concussed athletes. Extending this to injury prevention, Mojahed *et al*. [18] developed a nonlinear reduced-order model of the corpus callosum, which was used to evaluate the helmet protective performance [19]. Together, these multidisciplinary insights establish the corpus callosum as a critical structure for deciphering injury mechanisms, identifying neuroimaging biomarkers, and evaluating protective strategies.

Anatomically, the corpus callosum is commonly divided into three major subregions along the anteroposterior axis: the genu, midbody, and splenium, with the splenium being particularly vulnerable to traumatic injury. Postmortem neuropathological studies in humans have consistently identified the posterior corpus callosum as a frequent site of axonal injury [5-7, 20-23]. This regional vulnerability is strongly corroborated by *in vivo* neuroradiological findings, with abnormalities frequently detected in the splenium across multiple imaging modalities, including conventional computed tomography and magnetic resonance imaging (MRI) [24, 25], T2w spin echo imaging [26], proton magnetic resonance spectroscopy [27], and DTI [28-30]. More recently, Flusund *et al*. [31] constructed lesion frequency distribution maps from multimodal MRI data acquired in 269 patients with moderate-to-severe traumatic brain injury and found a preferential localization of traumatic axonal injury within the posterior half of the corpus callosum (Fig 1B). Despite the consistent clinical, neuropathological, and radiological evidence for this region-specific vulnerability, the biomechanical mechanisms underlying the heightened susceptibility of the splenium remain poorly understood.

**Fig 1.**
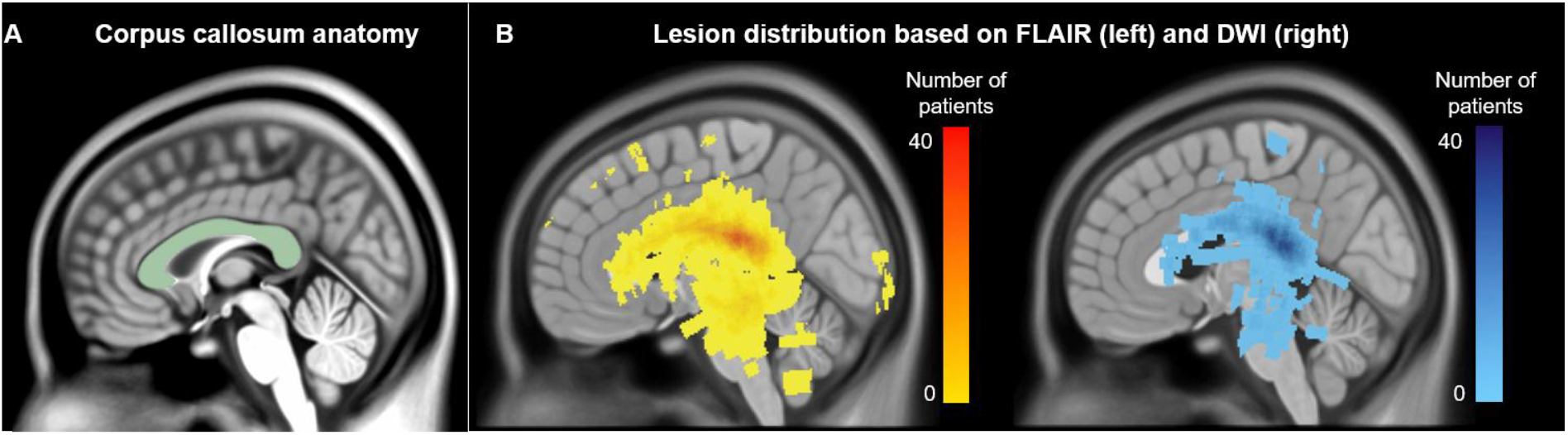
(A) Anatomy of the corpus callosum (highlighted in pale green) in the mid-sagittal plane. (B) Lesion frequency distribution maps of traumatic axonal injury on fluid-attenuated inversed recovery (FLAIR) and diffusion weighted imaging (DWI), both demonstrating a higher lesion frequency in the posterior corpus callosum (B). Images in subfigure B are adapted from Flusund *et al*. [31].

The falx cerebri has long been proposed as a contributing factor to the selective vulnerability of the splenium. Due to its anatomical proximity to the posterior corpus callosum and its substantially greater stiffness relative to brain tissue, the falx is hypothesized to amplify the mechanical loading and subsequent injury susceptibility within the splenium [24-26, 28]. However, direct verification of this hypothesis appears lacking. Previous biomechanical studies investigating the influence of the falx often treated the corpus callosum as a single structure [32-40]. Among the handful of FE head models with explicit representation of corpus callosum subregions [41-44], none have been used to isolate and quantify the specific mechanical influence of the falx on localized splenium responses.

This study aimed to uncover the mechanism of the selective vulnerability of the splenium by testing the hypothesis that the falx instigated amplified mechanical responses within the posterior subregion of the corpus callosum. Two FE models with and without an anatomically and mechanically detailed representation of the falx were used to simulate head impacts. By comparing the response of injury-related parameters across the corpus callosum subregions, the biomechanical role of the falx in driving splenial vulnerability was quantitatively clarified.

## Methods

### Finite element head model

The FE head model used in the current study, i.e., the KTH-Detailed head model (Fig 2), was previously developed by Zhou *et al*. [45] using LS-DYNA at the KTH Royal Institute of Technology. This model consisted of 4.2 million hexahedral elements and 0.5 million shell elements. The major head components were represented in the model, including the skull, brain, subarachnoid cerebrospinal fluid, ventricle, pia mater, dura mater, and tentorium. Its responses showed good correlation with brain-skull relative motion and brain strain [46, 47].

**Fig 2.**
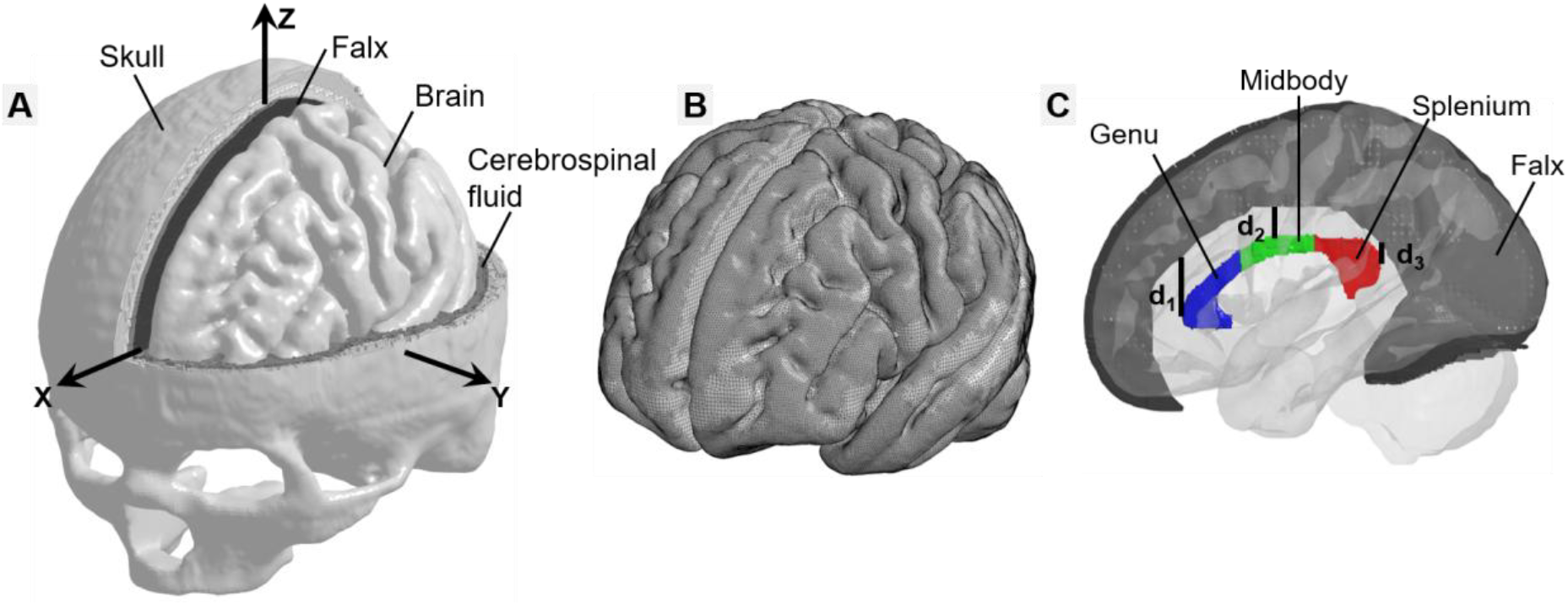
Finite element model of human head with detailed corpus callosum subregions. (**A**) Isometric view of the finite element head model with the skull open to expose the cerebrospinal fluid, brain, and falx. A skull-fixed coordinate system and corresponding axes are illustrated with the origin at the head’s centre of gravity. (**B**) Isometric view of the brain with visible meshlines. (**C**) Side view of the falx and three corpus callosum subregions (i.e., genu, midbody, splenium). The interspace distances between the lower edge of the falx and corpus callosum at the anterior, middle, and posterior sites are labelled at d1, d2, and d3, respectively.

Given that the current study focused on the influence of the falx on the response of corpus callosum subregions, the brain elements were further classified with reference to the ICBM-DTI-81 WM atlas [48] to have explicit representation of genu, midbody, and splenium (Fig 2C). The falx was modelled as membrane elements [49] with a thickness of 0.91 mm [50]. Similar to the approach of Lu *et al*. [34] and Alshareef *et al*. [51], the geometric profile of the falx was defined based on the segmentation from an open-access dataset [52], in which the falx had been automatically reconstructed using a fast-marching, multi-atlas-based segmentation from T1-weighted magnetic resonance imaging and susceptibility-weighted imaging [53]. As shown in Fig 2C, the model captured the non-uniform spatial relationship between the falx and the corpus callosum subregions, with the inferior edge of the falx located progressively closer to the splenium than to the genu and midbody. Following the anatomical measurements reported by Kayalioglu *et al*. [54], the interspace distances between the inferior edge of the falx and the corpus callosum were 18 mm anteriorly (d_1_), 10 mm centrally (d_2_), and 3.7 mm posteriorly (d_3_) (Fig 2C). These values fall within the range reported from cadaveric measurements (14.1 ± 5.6 mm at d_1_, 12.4 ± 4.7 mm at d_2_, and 2.1 ± 1.8 mm at d_3_), supporting the anatomical accuracy of the model geometry.

### Material properties

The material properties of each model component are listed in Table 1. The brain was modelled as an isotropic and homogeneous structure due to the lack of consensus regarding the mechanical anisotropy and heterogeneity of the brain [55, 56]. Specifically, a second-order Ogden-based hyperelastic constitutive law (Table 1B) was used to describe the nonlinear behavior of brain tissue, with six additional Prony-series linear viscoelastic terms (Table 1C) to account for its rate-dependent behavior [15]. The material properties of the falx/tentorium/dura mater were modeled as simplified rubber with the stress-strain curve obtained from the tissue experiment [57]. For the brain, its long-term shear modulus was around 1.05 kPa (calculated from the Ogden parameters in Table 1B based on the equation *G* = 0.5∑ *µ*_*i*_ · *α*_*i*_). For the falx, when fitting the linear region of the stress-strain curves, a Young’s modulus of around 31.5 MPa was attained. A direct comparison of the corpus callosum instantaneous hyper-elastic stress and falx stress in tension is shown in Fig 3, of which the falx was around 3 orders stiffer than the corpus callosum. In addition, the skull was modeled as rigid and separated the cortical bone from the diploe bone. The cerebrospinal fluid was modeled as an elastic fluid constitutive model and shared interfacial nodes with the brain and skull. The material behavior of the pia mater (thickness: 0.1 mm) was determined by the averaged material stress-strain curve from tissue experiments [58, 59].

**Table 1.** Material properties for the finite element head model (**A**) with Ogden-based hyperelastic material constants to describe the brain nonlinearity (**B**) and six Prony-series linear viscoelastic terms to capture the brain viscoelasticity (**C**). In Table A, K is the bulk modulus, and N/A is not applicable. In Table B, μ_i_ and α_i_ are Ogden parameters. In Table C, G_i_ and β_i_ represent the 6 shear relaxation moduli and corresponding decay constants.

| **A** | **Tissue** | **Young's modulus** | **Density (kg/dm<sup>3</sup>)** | **Poisson's ratio** | **Reference** |
| --- | --- | --- | --- | --- | --- |
|  | Cortical bone | 15 GPa | 2.00 | 0.22 | [15] |
|  | Diploe bone | 1 GPa | 1.30 | 0.24 |  |
|  | Brain | Hyper-Viscoelastic (Table 1 B-C) | 1.04 | ~0.5 |  |
|  | Cerebrospinal fluid | K = 2.1 GPa | 1.00 | N/A |  |
|  | Falx & Tentorium & Dura mater | Average stress-strain curve (Fig 2) | 1.13 | N/A | [57] |
|  | Pia mater | Average stress-strain curve | 1.13 | N/A | [58] |
| **B** | **Ogden parameter** | **Value** | **Ogden parameter** | **Value** | **Reference** |
| | $\mu_1$ (Pa) | 53.8 | $\alpha_1$ | 10.1 | [15] |
| | $\mu_2$ (Pa) | -120.4 | $\alpha_2$ | -12.9 | |
| **C** | **Shear relaxation moduli** | **Value** | **Decay constant** | **Value** | **Reference** |
| | $G_1$ (MPa) | 0.32 | $\beta_1$ (s<sup>-1</sup>) | 10<sup>6</sup> | [15] |
| | $G_2$ (kPa) | 78 | $\beta_2$ (s<sup>-1</sup>) | 10<sup>5</sup> | |
| | $G_3$ (kPa) | 6.2 | $\beta_3$ (s<sup>-1</sup>) | 10<sup>4</sup> | |
| | $G_4$ (kPa) | 8.0 | $\beta_4$ (s<sup>-1</sup>) | 10<sup>3</sup> | |
| | $G_5$ (kPa) | 1.0 | $\beta_5$ (s<sup>-1</sup>) | 10<sup>2</sup> | |
| | $G_6$ (kPa) | 3.0 | $\beta_6$ (s<sup>-1</sup>) | 10<sup>1</sup> | |

**Fig 3.**
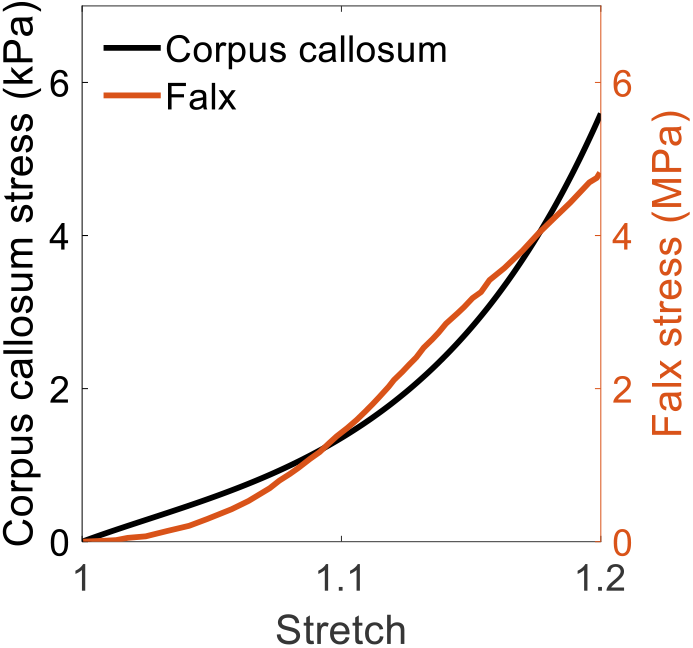
Comparison of the hyper-elastic stress of the corpus callosum (solely based on the Ogden parameters in Table 1B) and stress-stretch curve of the falx in tension. Because the stress magnitudes differed between the corpus callosum and falx, separate y-axes were used to facilitate comparison, with corpus callosum stress shown on the left axis (in black) and falx stress on the right axis (in orange).

### Impact loadings

To study the influence of the falx on the corpus callosum subregion, whole-head impact simulations were performed. Ten cases, covering different loading directions and severities, were selected from a National Football League impact dataset, in which head kinematics were obtained from laboratory reconstructions of on-field video-recorded collisions using Hybrid III anthropometric test dummies [60]. The acceleration peaks of selected cases are summarized in Table 2, and detailed acceleration profiles of one representative case (i.e., Case 1) are exemplified in Fig. 4. For each impact, the directional linear acceleration and angular velocity curves were imposed to one node located at the head’s center of gravity and constrained to the rigid skull. Each impact was simulated using the head model with and without the falx, resulting in a total of 20 simulations. For an impact of 50 ms, it took 13.45 hours for the model with falx, and 13.25 hours for the model without falx, solved by the massively parallel processing version of LS-DYNA 15.0 version with double precision using 128 central processing units.

**Table 2.** Peaks of linear and angular accelerations of the 10 simulated cases. Note that the X, Y, and Z axes are the same as those in the skull-fixed coordinate system in Fig 1A.

| Case ID | Peak linear acceleration (g) |  |  |  | Peak angular acceleration (krad/s<sup>2</sup>) |  |  |  |
| --- | --- | --- | --- | --- | --- | --- | --- | --- |
|  | X | Y | Z | Magnitude | X | Y | Z | Magnitude |
| Case 1 | -78.0 | 66.5 | -21.8 | 102.1 | -3.21 | -2.39 | 4.40 | 4.98 |
| Case 2 | -29.2 | 22.5 | -34.6 | 49.4 | -2.38 | -1.01 | 1.10 | 2.77 |
| Case 3 | 19.7 | 56.3 | -6.0 | 59.8 | -4.04 | 2.95 | -6.74 | 8.30 |
| Case 4 | -21.5 | 59.4 | -57.8 | 78.8 | -5.82 | 1.66 | 2.43 | 6.23 |
| Case 5 | -31.9 | -133.3 | -41.6 | 138.2 | 4.65 | 1.35 | 6.82 | 7.50 |
| Case 6 | -42.3 | 114.3 | -27.4 | 123.7 | -3.53 | -1.70 | 5.53 | 6.37 |
| Case 7 | -64.1 | 79.9 | 44.5 | 103.2 | -5.69 | -4.70 | 3.88 | 7.54 |
| Case 8 | -31.7 | 40.0 | -18.2 | 52.3 | -2.77 | -1.87 | -2.21 | 3.13 |
| Case 9 | -62.3 | 54.9 | 29.9 | 86.5 | -2.35 | -1.57 | 3.15 | 3.98 |
| Case 10 | 30.7 | -78.3 | 42.1 | 84.0 | 5.31 | 2.58 | 3.90 | 6.39 |

**Fig 4.**
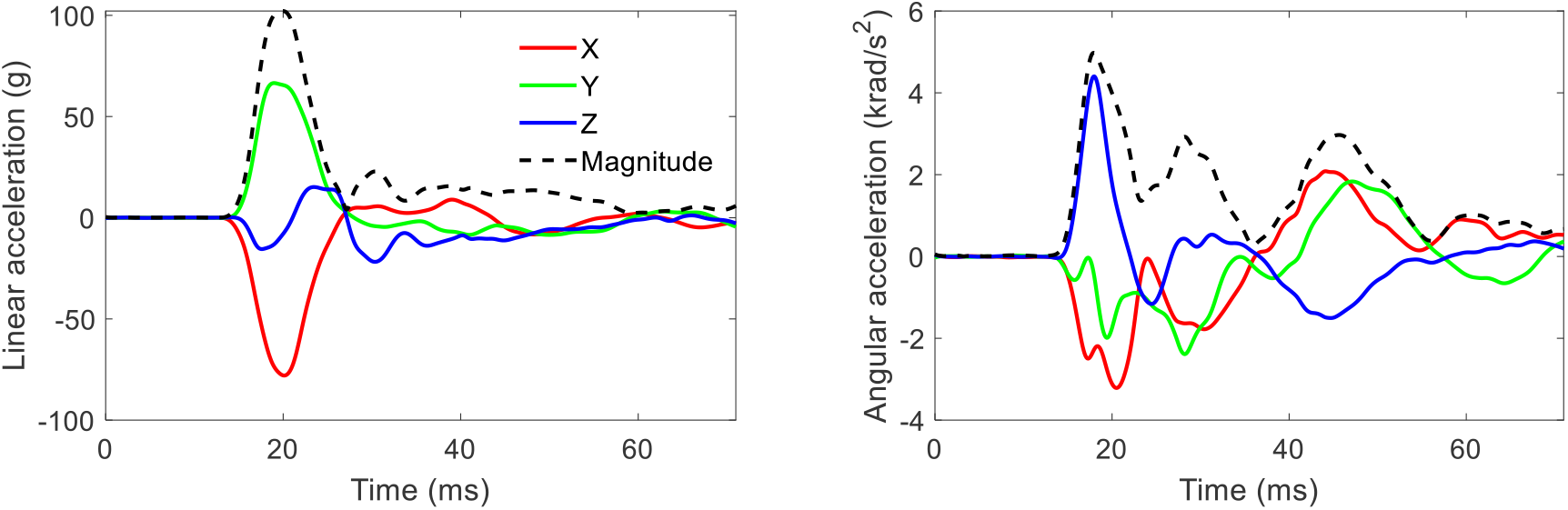
Time-history curves of linear acceleration (left) and angular acceleration (right) of Case 1. Note that the X, Y, and Z axes are the same as those in the skull-fixed coordinate system in Fig 2A.

### Data analysis

In each simulated impact, the mechanical responses of the genu, midbody, and splenium were extracted. The strain and strain rate were employed to quantify the tissue deformation and calculated as the first principal values of the Green-Lagrange strain tensor and the rate of deformation tensor, respectively, at each time step [61, 62]. Given that the brain is particularly vulnerable to deformation secondary to the shear force [63, 64], the maximum shear stress was used to characterize the localized tissue force. At the element level, peak values of strain, strain rate, and shear stress across the entire impact duration were used as injury metrics. At each corpus callosum subregion, the peak strain, strain rate, and shear stress values of all affiliated elements were calculated. To avoid potential numerical artefacts, 95^th^ percentile peak values were reported for each corpus callosum subregion [65, 66].

The effects of the falx on the subregions of the corpus callosum were assessed separately for peak strain, peak strain rate, and peak shear stress using linear mixed-effects models, with falx condition (with vs. without) and corpus callosum subregion (genu, midbody, splenium) included as fixed effects, and impact case as a random intercept to account for repeated measurements across impact scenarios. To test the hypothesis that the falx preferentially instigated larger mechanical loading in the splenium, paired comparisons were performed on the falx-induced changes (the difference with and without falx), comparing the splenium with the genu and the midbody. The threshold for statistical significance was set at *p* < 0.05.

## Results

### Strain and strain rate in the corpus callosum subregions

We first aimed to elucidate the changes in strain and strain rate in the corpus callosum subregions due to the presence of the falx. For one illustrative impact (Case 1), the inclusion of the falx was associated with relatively greater strain and strain rate in the splenium. As qualitatively shown in Fig 5A-B, when the falx was presented, regions of high strain and strain rate (indicated by the red areas) were primarily located in the posterior corpus callosum, corresponding to the splenium. In contrast, when the falx was omitted, the magnitude of strain and strain rate in the corpus callosum were generally reduced, with relatively higher responses shifting toward the anterior region (i.e., the genu). These qualitative observations were further supported by quantitative analysis of peak strain and strain rate in three corpus callosum subregions (Fig 5C-D). With the falx included, the splenium exhibited the highest peak strain (0.32), compared with 0.25 in both the genu and midbody. Similarly, the highest peak strain rate was in the splenium (30.1 /s), followed by the genu (26.8 /s) and midbody (19.9 /s). In the absence of the falx, the highest peak strain and strain rate were observed in the genu (0.27 and 25.9 /s, respectively), whereas the lowest values were found in the midbody (0.15 and 14.9 /s, respectively) and the intermediate values in the splenium (0.21 and 16.2 /s, respectively).

**Fig 5.**
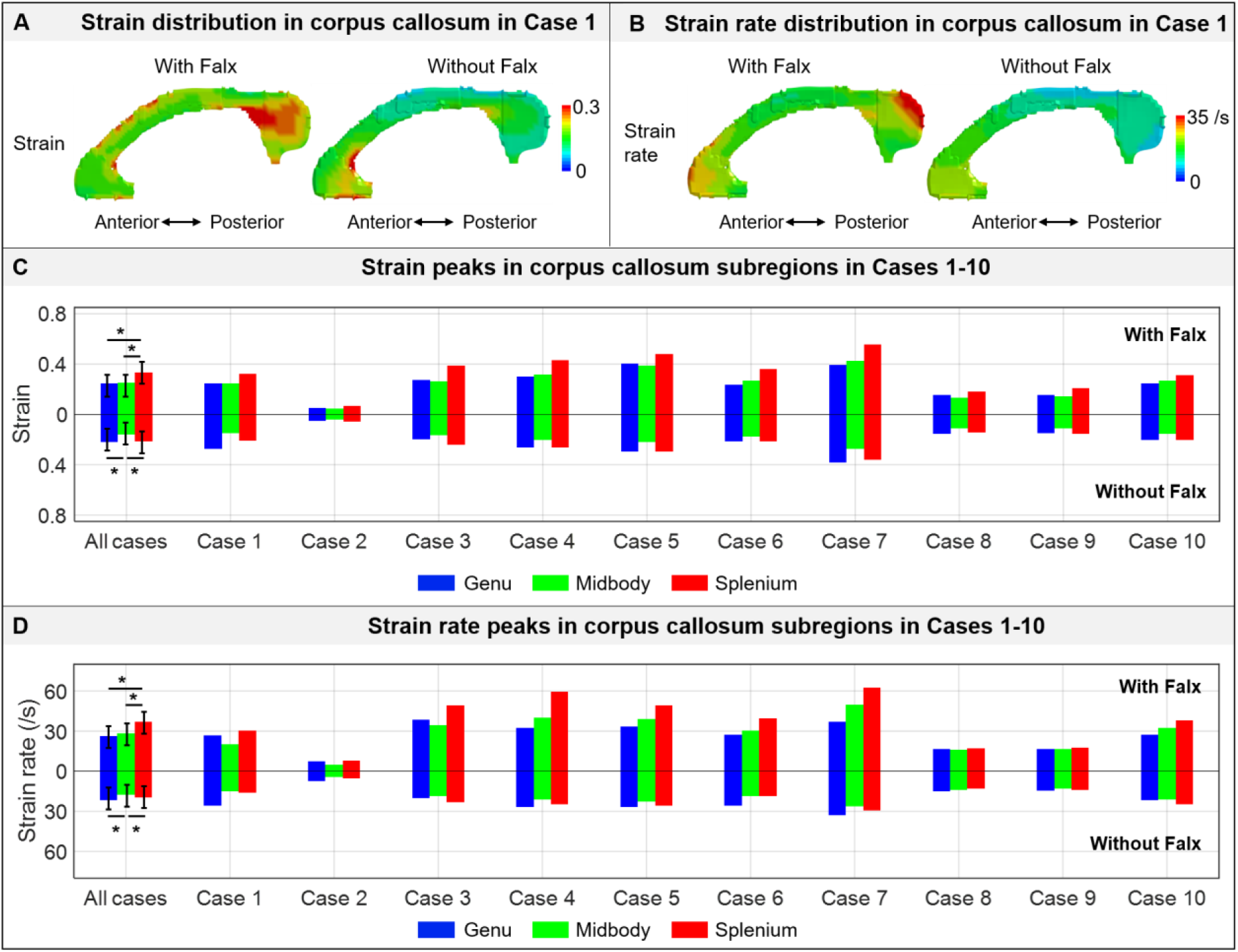
Influence of the falx on the strain and strain rate response in the corpus callosum subregions (i.e., genu, midbody, and splenium). (**A**) Strain distribution in the corpus callosum in Case 1. (**B**) Strain rate distribution in the corpus callosum in Case 1. (**C**) Strain peak in three corpus callosum subregions in Cases 1-10. (**D**) Strain rate peak in three corpus callosum subregions in Cases 1-10. Note that, in subfigures A and B, the anterior and posterior direction of the corpus callosum is indicated. In subfigures C and D, the upper panel shows the results from simulations with the falx, while the lower panel without the falx. * indicates the difference in the referred pair is statistically significant.

At the group level, encompassing all 10 simulated impacts, we compared the peak strain and peak strain rate in the three corpus callosum subregions (Fig 5C-D). Across all simulated impacts, inclusion of the falx consistently resulted in higher peak strain (upper panel, Fig. 5C) and peak strain rate (upper panel, Fig. 5D) in the splenium than in the genu and midbody. When the falx was included, the splenium exhibited the highest peak strain (mean ± standard deviation, 0.33 ± 0.15), followed by the midbody (0.25 ± 0.12) and genu (0.25 ± 0.11). Similarly, the splenium showed the highest peak strain rate (37.1 ± 18.6 /s), followed by the midbody (28.3 ± 13.6/s) and genu (26.2 ± 10.0 /s). In the absence of the falx, the highest peak strain and peak strain rate were observed in the genu (0.22 ± 0.09 and 25.9 ± 7.6 /s, respectively), whereas the lowest values were found in the midbody (0.16 ± 0.07 and 17.5 ± 6.3 /s, respectively) and the intermediate values in the splenium (0.21 ± 0.09 and 19.5 ± 7.3 /s, respectively).

When we statistically analysed the peak strain across the 10 simulated cases, the linear mixed-effects model showed no significant overall main effect of falx condition (*p* = 0.067), but a significant interaction between falx condition and corpus callosum subregion, indicating region-dependent effects of the falx. Paired comparisons showed that the falx-induced increase in peak strain was significantly larger in the splenium than in the genu (*p* < 0.001) and the midbody (*p* = 0.006). For peak strain rate, no significant overall main effect of falx condition was observed (*p* = 0.063), whereas the falx exerted a significant region-specific effect in the splenium (interaction *p* < 0.001). The falx-induced increase in strain rate was significantly greater in the splenium than in the genu (*p* = 0.003) and midbody (*p* = 0.002).

### Stress in the corpus callosum subregions

We next examined the shear stress responses in the corpus callosum subregions to explain the mechanistic cue of relatively more severe deformation observed in the splenium. When the falx was included, the splenium consistently exhibited the highest peak shear stress among the three corpus callosum subregions across all simulated impacts (Fig. 6A, upper panel). The peak shear stress (mean ± standard deviation) was 3.93 ± 2.30 kPa in the splenium, compared to 2.88 ± 1.66 kPa in the midbody and 2.46 ± 1.23 kPa in the genu. In contrast, when the falx was absent (Fig 6A, lower panel), the genu exhibited the highest peak shear stress (2.46 ± 1.27 kPa), followed by the splenium (2.42 ± 1.16 kPa) and the midbody (1.88 ± 0.88 kPa). Element-wise distributions of peak shear stress for two representative cases are qualitatively shown in Fig. 6B (Case 1) and Fig. 6C (Case 7). In both cases, inclusion of the falx resulted in markedly elevated shear stress concentrations in the splenium, whereas omission of the falx shifted the relatively higher stress concentrations toward the genu.

**Fig 6.**
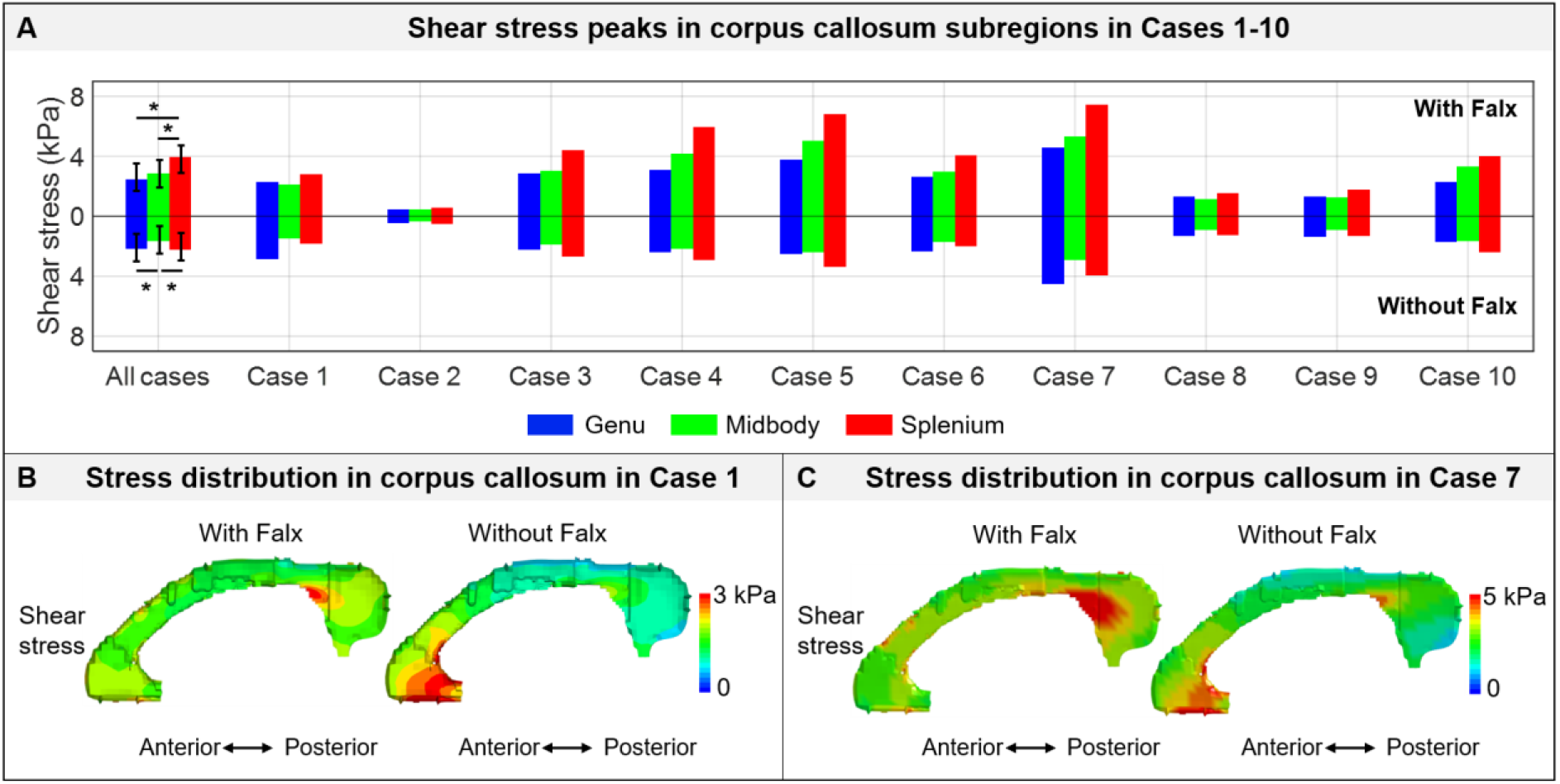
Influence of the falx on the stress response in the corpus callosum subregions (i.e., genu, midbody, and splenium). (**A**) Strain peak in three corpus callosum subregions in Cases 1-10, with the falx-included results in the upper panel and falx-excluded results in the lower panel. * indicates the difference in the referred pair is statistically significant. (**B**) Strain distribution in the corpus callosum in Case 1. (**C**) Strain distribution in the corpus callosum in Case 7.

We statistically analysed the peak shear stress in the 10 simulated impacts and found a significant dependence of the falx effect on corpus callosum subregion, as indicated by the significant falx-subregion interaction in the linear mixed-effects model (*p* = 0.001), whereas the overall main effect of falx condition was not significant (*p* = 0.330). Subsequent paired comparisons demonstrated that the stress increase associated with the presence of the falx was more pronounced in the splenium than in the genu (*p* = 0.002) and midbody (*p* = 0.006). A greater falx-related stress increase was also observed in the midbody compared with the genu (*p* = 0.002).

## Discussion

The current study evaluated the long-standing clinical hypothesis that the falx cerebri contributes to the selective vulnerability of the splenium within the corpus callosum. We systematically compared impact-induced responses estimated by high-resolution FE head models - with and without a detailed falx representation. The result demonstrated that the falx selectively increased the mechanical loading within the splenium, characterized by significantly elevated splenial strains, strain rates, and shear stresses relative to the genu and midbody. This work provided a plausible mechanistic explanation for the frequent clinical observation of splenium involvement in TBI patients.

Our computational results directly suggested that the splenial susceptibility was attributed to the coupled effects of the anatomical configuration of the falx relative to the corpus callosum and the marked stiffness mismatch between these structures. Anatomically, the falx extends progressively deeper into the cranial cavity toward the posterior region, positioning its inferior edge in closest proximity to the corpus callosum near the splenium (Fig 2C). Mechanically, the falx is several orders of magnitude stiffer than the corpus callosum (e.g., an approximated elastic modulus of ∼31.5 MPa for the falx vs. a long-term shear modulus of 1.05 kPa for the corpus callosum) (Fig 3). Consequently, the falx’s influence on corpus callosum mechanics is not uniform along the anteroposterior axis. In the anterior region, less doze of forces is transferred to the corpus callosum with relatively smaller strain and strain rate in the genu. In the posterior region, the falx is elongated and effectively transmits a larger amount of loading to the corpus callosum, resulting in more severe deformation in the splenium. This proposed mechanism is supported by the observed redistribution of shear stress (Fig. 6), in which inclusion of the falx consistently increased strain and strain rate most prominently in the splenium (Fig. 5) in the current study.

Indirect support for the proposed role of the falx in splenial vulnerability came from experimental animal models, in which regional patterns of axonal injury varied with species-specific intracranial anatomy. Several studies from the University of Pennsylvania [67-69] characterized the brain pathology in the porcine following inertial closed head injury and reported that the corpus callosum, especially the splenium, was not damaged to nearly the same extent as typically found in brain-injured humans. [69] suggested that this pathological disparity might be related to the species-specific anatomical differences, as the porcine falx extends only superficially between the hemispheres and therefore likely exerts less biomechanical influence on the corpus callosum than the human falx. Although direct cross-species biomechanical comparisons should be interpreted with caution, these observations were broadly consistent with the mechanism proposed in the current study and further supported the importance of the falx in instigating preferentially greater splenial deformation in the human corpus callosum.

Several previous computational studies using alternative FE head models investigated the biomechanical responses of corpus callosum subregions based on the maximum principal strain (as was also used herein), although the selective amplification of splenial strain was not consistently reported. For example, Ji *et al*. [70] employed one FE model (falx material: linear elastic with a Young’s moduli of 31.5 MPa; falx thickness: not reported) to simulate 477 sports-related head impacts from three independent datasets and reported that the maximum principal strain was consistently concentrated in the central portion of the corpus callosum, approximately corresponding to the midbody region in the present anatomical partitioning. Montanino *et al*. [71] used another FE head model (falx material: same as the nonlinear stress-strain curve in Fig 2; falx thickness: 1.5 mm) to simulate one concussive impact and found the maximum corpus callosum strain was located in the midbody. Although the exact reasons behind these cross-model differences in the spatial distribution of maximum principal strain within corpus callosum subregions remain unclear, differences in falx modelling assumptions (e.g., falx geometry and its anatomical relationship to the corpus callosum) might contribute to the observed discrepancies.

Our results demonstrated that the mechanical influence of the falx within the corpus callosum was highly heterogeneous, rather than acting as a uniform amplifier of tissue deformation throughout the entire structure. As shown in Fig 4, the presence of the falx consistently increased the strain in the splenium and, to a lesser extent, the midbody, whereas this amplifying trend was not consistently observed in the genu. In contrast, previous FE studies, such as Ho *et al*. [33] and Hernandez *et al*. [43], often evaluated the influence of the falx by treating the callosal structure as a single structure. The current results suggest that averaging deformation over the entire corpus callosum might obscure region-specific effects induced by the falx within the corpus callosum. It should be noted that the heterogeneous effect of the falx has been previously reported with the region of interest across the whole brain [34, 37, 38].

The current study modelled the corpus callosum as an isotropic and homogeneous material, whereas a body of literature reported that the corpus callosum exhibited subregional heterogeneity in terms of material properties and microstructure representation. For example, under *in vivo* conditions, Johnson *et al*. [72] found that the genu had significantly lower storage moduli and higher loss moduli than the midbody, while Smith *et al*. [73] reported that the tensile anisotropy of the genu was significantly lower than that of the midbody and splenium. Under *in vitro* conditions, Reiter *et al*. [74] reported pronounced anisotropic mechanical behaviour in the corpus callosum based on cyclic compression-tension and simple shear test, although an early study from the same group found no significant directional dependence when the tissue sample from the corpus callosum was tested along and perpendicular to the fiber direction [75]. In addition to mechanical heterogeneity, substantial regional variations in axonal composition have also been reported. For example, Aboitiz *et al*. [76] showed that thin fibers were most densely distributed in the genu, decreased toward the posterior midbody, and increased again in the splenium. Initial attempts have been made to integrate such microstructural information into a multi-scale framework to investigate the load transmission from the whole head level (where the impact occurred) to the molecular level (where the injury occurred) [71]. How these regional differences in constitutive behaviour and fiber architecture contribute to the vulnerability of the splenium warrants future investigations.

## Conclusion

This study aimed to answer why the splenium is so commonly affected by traumatic impacts by comparing responses in corpus callosum subregions estimated by paired high-resolution FE head models with and without an anatomically and mechanically detailed representation of the falx. Our results showed that the presence of the falx consistently increased strain, strain rate, and shear stress in the splenium to a greater extent than in the genu and midbody, verifying the hypothesis that the mechanically stiffer falx with anatomical proximity to the posterior of the corpus callosum contributes to the selective vulnerability of the splenium. Our work provided a plausible explanation of the prevalence of the splenial injury in TBI patients and yielded new knowledge on the heterogeneous effect of the falx on brain subregions.

## Acknowledgments

This research has received funding from KTH Royal Institute of Technology (Stockholm, Sweden), Swedish Research Council (VR-2024-05848, and VR-2024-02782). The content of this article is solely the responsibility of the authors and does not necessarily represent the official views of neither funding agencies. The computational simulations were enabled by resources provided by the National Academic Infrastructure for Supercomputing in Sweden (NAISS) at the center for High Performance Computing (PDC) partially funded by the Swedish Research Council through grant agreement no. 2026/3-281 and 2026/4-723.

